# Characterization of the *Plasmodium falciparum* homologue of Vps16, a member of the Vps-C tethering complex

**DOI:** 10.1101/2024.10.07.617026

**Authors:** Thomas Galaup, Florian Lauruol, Audrey Sergerie, Dave Richard

**Affiliations:** Centre de Recherche en Infectiologie, CRCHU de Québec-Université Laval, 2705 Boul. Laurier Québec (QC), Canada, G1V 4G2; Department of Microbiology-Infectious Diseases and Immunology, Faculty of Medicine, Laval University. Quebec City (Qc), Canada

**Author notes:** Experimental Hematology, University Medical Center Groningen, Hanzeplein 1, 9700 RB, DA13, Groningen, The Netherlands. Corresponding author and Lead Contact: (DR).

**Keywords:** Malaria, protein trafficking, rhoptries, tethering complex

## Abstract

The organelles of the apical complex (rhoptries, micronemes and dense granules) are critical for erythrocyte invasion by the malaria parasite *Plasmodium falciparum*. Though they have essential roles in the parasite lifecycle, the mechanisms behind their biogenesis are still poorly defined. The Class C Vps proteins Vps11, 16, 18 and 33 constitute the core of the CORVET and HOPS complexes implicated in vesicle tethering and fusion in the eukaryotic endolysosomal system. Work in the model apicomplexan *Toxoplasma gondii* has revealed that TgVps11 is essential for the generation of the apical complex. *P. falciparum* possesses all the four subunits of the Vps-C complex but their function is currently unknown. We here present an initial characterization of the *P. falciparum* orthologue of Vps16, a member of the Vps-C complex. Our structural predictions suggest that the structure of PfVps16 is similar to its other eukaryotic counterparts and that the binding region responsible for its interaction with PfVps33 is conserved. We next show that PfVps16 is expressed throughout the asexual erythrocytic cycle and that it is potentially associated with the rhoptries in schizont stage parasites. Finally, we present our unsuccessful attempts at studying its function using knock sideways.

## INTRODUCTION

Although significant strides have been made in decreasing the death and illness rates associated with malaria in recent years, the disease continues to impose a substantial burden on various tropical and sub-tropical regions worldwide. In 2022 alone, an estimated 249 million cases causing over 608,000 deaths were attributed to *Plasmodium* parasites, with a majority caused by *P. falciparum*, the species responsible for the most severe form of the disease(1). The emergence of resistance to many existing antimalarial drugs, including the primary treatment artemisinin(2, 3), underscores the urgent necessity for the development of new intervention approaches.

*Plasmodium* species are obligate intracellular parasites that begin their lifecycle by invading host cells. The invasion of erythrocytes by malaria merozoites involves a complex, multistep process driven by the sequential release of organelles that make up the apical complex: rhoptries, micronemes, and dense granules(4). These organelles are newly formed during a unique cell division process called schizogony(5, 6). Work on intracellular protein trafficking in the model apicomplexan *T. gondii* has led to the idea that its endosomal system has been evolutionarily repurposed for rhoptry and microneme formation(7, 8). The mechanisms behind the formation of the apical complex in *P. falciparum* are not as well understood. Conditional knockdown of the Golgi-resident escort protein PfSortilin(9, 10) has shown that it is required for the trafficking of proteins to the micronemes, rhoptries and dense granules(11). The absence of the Rhoptry apical membrane antigen (RAMA) leads to dysmorphic rhoptry necks highlighting its importance in their proper biogenesis(12). The Cytosolically exposed rhoptry-leaflet-interacting proteins 1 and 2 (CERLI1 and 2) localize to the outside of the rhoptry membrane and their absence alters the distribution of rhoptry proteins indicating that they are likely to also play a role in their formation(13-15). In *T. gondii*, an intermediate endosome-like compartment (ELC) is involved in protein sorting between the Golgi apparatus and the apical organelles(16) but evidence of a direct pathway from the Golgi apparatus in *P. falciparum*, suggests that the ELC may not be necessary in this organism(9). Partial colocalization of the *P. falciparum* homologues of the small G-protein Rab11A and Adaptor protein 1 with rhoptry markers have led to the suggestion that these proteins might play a role in the process of vesicular fusion at the rhoptry membrane(17, 18).

Eukaryotic cells possess several tethering complexes that capture intracellular vesicles to initiate fusion to their target membrane(19). These complexes are found at distinct subcellular locations and are key to providing specificity(20, 21). CORVET (class C core vacuole/endosome tethering) and HOPS (homotypic fusion and vacuole protein sorting) are tethering complexes associated with the early or late endosomes, respectively, where they play critical roles in processes such as the biogenesis of lysosomes and autophagy(22). They share a conserved core of four proteins, Vps11, Vps16, Vps18 and Vps33, referred to as the Vps-C core(23) and, in yeast and mammalian cells, two specific subunits each (Vps3 and 8 for CORVET; Vps39 and 41 for HOPS). The Vps-C core of both CORVET and HOPS share strong structural homology whilst the specific subunits are markedly different(24). Intriguingly, the Vps-C core proteins are found in *P. falciparum* however, only the CORVET-specific Vps3 is conserved and no Vps8 or either of the HOPS-specific subunits are detectable(23, 25) suggesting specific adaptations of its endolysosomal system. In *T. gondii*, conditional knock-down of TgVps11 and TgVps18 led to the disruption of the biogenesis of the rhoptries, dense granules and a subset of micronemes(26).

In this manuscript, we present an initial characterization of the Vps-C core protein PfVps16.

## RESULTS AND DISCUSSION

### Analysis of the predicted 3D structure of PfVps16

We first examined the 3D structure prediction of PfVps16 generated by the AlphaFold3 algorithm(27). The protein exhibits a β-propeller structure in the regions from amino acids 1 to 173 and 340 to 481 and an ɑ-solenoid structure from amino acid 482 to the C-terminus (Fig. 1A). This domain architecture is also found in Vps-C proteins from yeast and humans and share similarity with components of vesicular coats and proteins found in the nuclear pore(22, 28) (Fig. 1B and C). Intriguingly, a region of structural uncertainty, indicated by a low predicted local-distance difference test (pLDDT) value, is found from amino acids 174 to 339 in the middle of the β-propeller structure (Fig. S1). The structural alignment of the predicted 3D structure of PfVps16 with the known 3D structure of *Saccharomyces cerevisiae* Vps16 resulted in a TM-Score of 0.72804 (Fig. 1D). This high TM-Score demonstrates significant structural similarity between the two proteins, apart from the insertion in the N-terminal domain of PfVps16. In human cells, the interaction of Vps33A with the core of the Vps-C complex formed by Vps-11 and 18 requires Vps16(29-32) and this is achieved through a helical fragment containing amino acids 642 to 736 of Vps16(33). Human Vps33A consists of three domains: domain 1, domain 2, and domain 3, which is further subdivided into 3a and 3b(33) (Fig. 2A). The AlphaFold prediction of PfVps33 (Fig. 2B and S2A) shows two structured regions with high pLDDT values (residues 1 to 213 and 726 to 1042) and a central disordered region with a low pLDDT value corresponding to residues 214 to 725 (Fig. S2A and B). The structural alignment of the predicted 3D structure of PfVps33 (Fig. 2C) with the known 3D structure of human Vps33A yielded a TM-Score of 0.88927 indicating important homology (Fig. 2C). In humans, Vps33A interacts with Vps16 through its domain 3b, specifically with amino acids 642-736 of Vps16(33) (Fig. 2D). By aligning these residues with PfVps16, we identified the corresponding region as amino acids 837 to 943 (Fig. S2C, TM-score = 0.87154). Using AlphaFold3, we then predicted the interaction between PfVps16 (837-943) and PfVps33, suggesting that PfVps16 interacts with the 3b domain of PfVps33 (ipTM = 0.84) (Fig. 2E). These interaction predictions suggest that PfVps33 and PfVps16 interact in a similar way to what has been observed in humans. However, unlike the human protein, PfVps33 appears to have low-complexity regions (LCRs) splitting domain 2 in two. LCRs are fairly common in *P. falciparum*(35) but their role remains debated. One of the LCRs in this region (residues 314 to 405) is composed of 68.48% aspartic acid (D) and 25% glutamic acid (E). It has been suggested that the D/E repeats may be involved in gene regulation(36).

**Figure 1:**
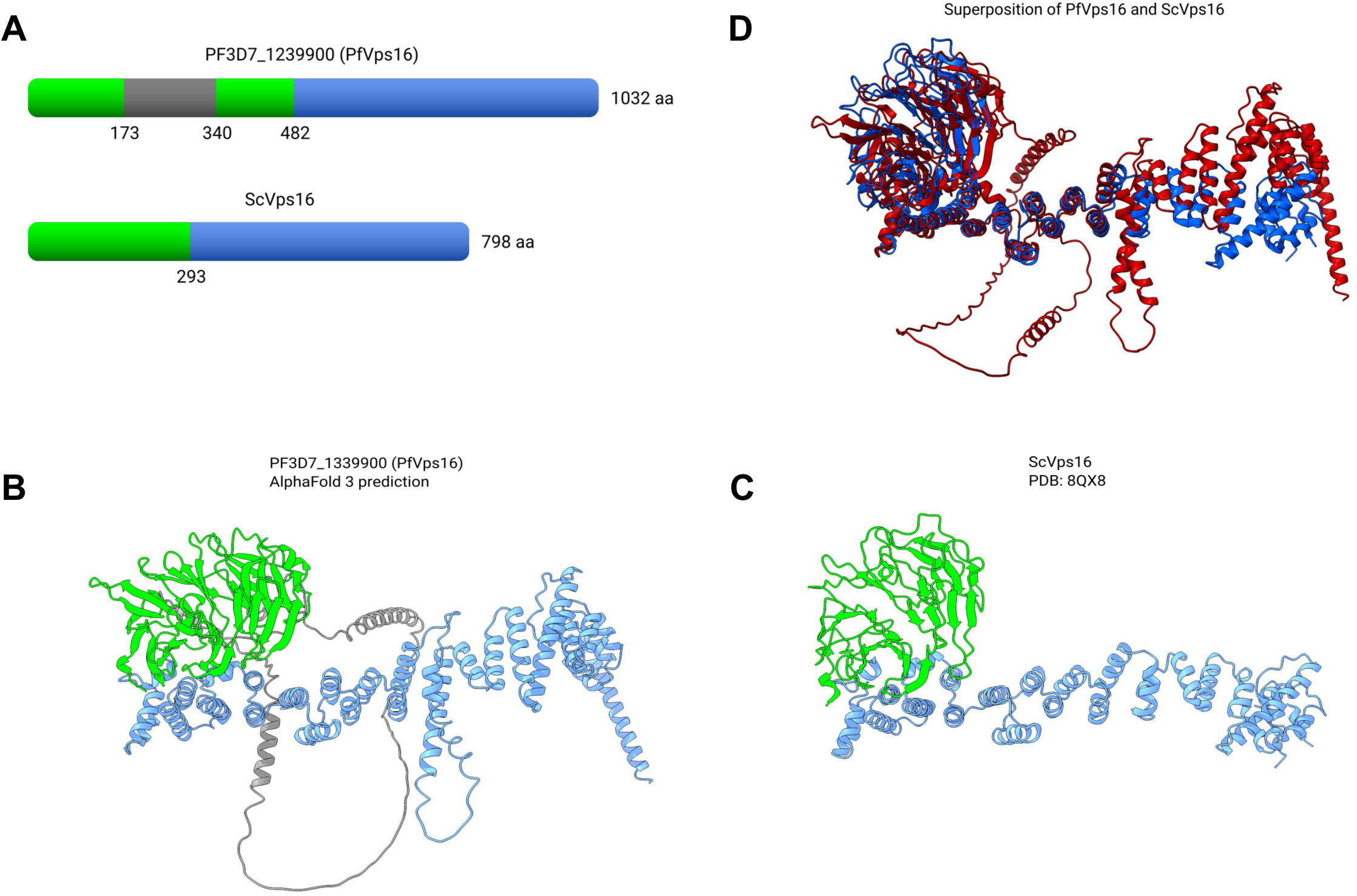
Structural prediction of PfVps16. (A) Predicted domains of PfVps16 and *S. cerevisiae* Vps16. Amino acids positions of β-propeller structures (green), α-solenoids (blue) and unpredicted structure (grey). (B) AlphaFold 3 structure prediction of PfVps16. Predicted β-propellers in green, predicted α-solenoids in blue, and disordered domain corresponding to low pLDDT value in grey. (C) *Sc*Vps16 structure obtained from experimental data (PDB: 8QX8) with β-propellers in green and α-solenoids in blue. (D) Superposition of PfVps16 in red and ScVps16 in blue. Superposition was obtained using the matchmaker tool from ChimeraX.

**Figure 2:**
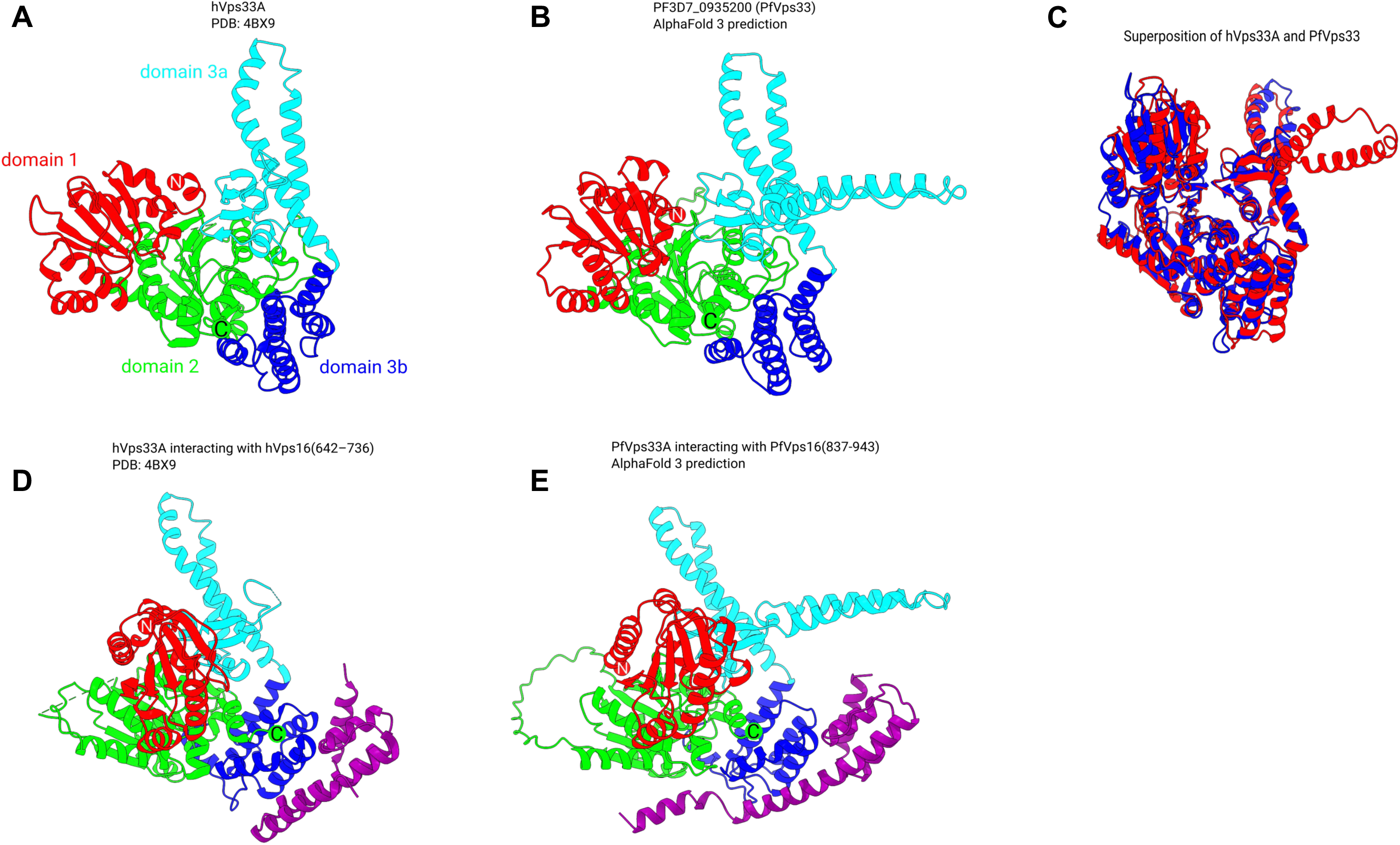
Structural prediction of PfVps33 and its interaction with PfVps16. (A) The structure of human Vps33A (hVps33A), obtained from experimental data (PDB: 4BX9), is shown with its three domains color-coded: domain 1 in red (residues 1 to 141), domain 2 in green (residues 142 to 242 and 473 to 595), domain 3a in cyan (residues 243 to 368), and domain 3b in blue (residues 369 to 472). (B) The AlphaFold 3 predicted structure of PfVps33 with the three domains similarly color-coded: domain 1 in red (residues 1 to 140), domain 2 in green (residues 141 to 213, 726 to 744 and 1019 to 1147), domain 3a in cyan (residues 745 to 908), and domain 3b in blue (residues 909 to 1018). Domain 2 is interrupted by a disordered region not shown in the figure (residues 214 to 725). (C) Superposition of hVps33 in blue and PfVps33 in red (disordered domain not shown). (D) The interaction of hVps33A with hVps16 (residues 642-736, in purple). (E) The predicted interaction of PfVps33 with PfVps16(residues 837-943, in purple).

### PfVps16 is expressed throughout the asexual erythrocytic cycle

To determine the subcellular localization of PfVps16, we tagged the endogenous gene at the 3’ end with GFP by single cross-over recombination using the selection-linked integration (SLI) strategy(37). To allow the functional analysis of PfVps16 by knock sideways (Ks, see below), a double FK506 binding protein domain (2xFKBP) tag was also included (PfVps16-2xFKBP-GFP, referred to as PfVps16Ks throughout the remainder of the manuscript)(37) (Fig. 3A). PCR analysis of genomic DNA from a clonal line revealed proper 5’ and 3’ integration of the plasmid at the Vps16 locus and the absence of a WT allele (Fig. 3B). Western blots on mixed-stages parasite extracts of the clonal line using an anti-GFP antibody revealed a specific single band at the expected size of around 170 kDa for the PfVps16Ks fusion protein. Antibodies against the constitutive protein HSP70 were used as loading controls (Fig. 3C). To determine the expression profile of PfVps16Ks throughout the erythrocytic cycle, we performed fluorescence microscopy on tightly synchronized parasites taken at different stages of the cycle. This revealed that the protein was detectable as a single focus in rings, one or more foci in trophozoites and multiple foci in schizonts (Fig. 3D).

**Fig. 3:**
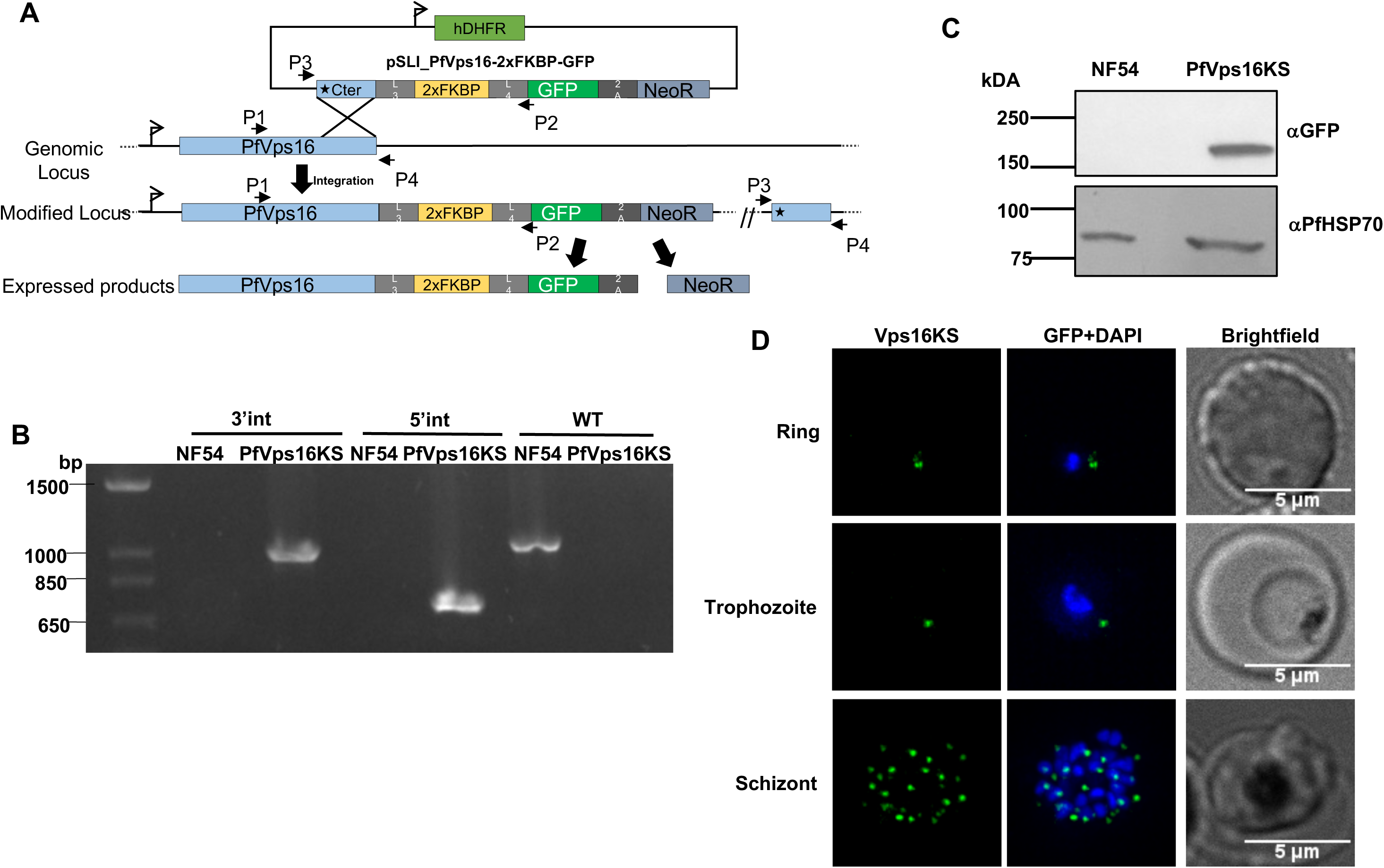
PfVps16 is expressed throughout the asexual erythrocytic cycle. (A) Schematic showing the tagging strategy by single cross-over recombination using SLI. (B) PCR on parasite genomic DNA showing the proper integration of the tagging vector at the PfVps16 locus (5’ junction: primers P1 and P2, 3’junction: primers P3 and P4) and the disappearance of the WT allele in the PfVps16Ks line (primers P1 and P4). (C) Western blot on parasite protein extracts showing the expression of PfVps16Ks at the expected size. PfHSP70 is used as a control. (D) PfVps16Ks is expressed throughout the asexual erythrocytic cycle. Scale bar represents 5μm. Blue: DAPI stained nucleus.

### Subcellular localization of PfVps16

We next looked at the subcellular distribution of PfVps16Ks in schizonts by immunofluorescence assays (IFA). The pattern of fluorescence being reminiscent of the behavior of proteins found at the Golgi apparatus(9, 10, 38-42) we first determined whether PfVps16Ks colocalized with the cis-Golgi marker ERD2. Interestingly, though the foci were often close, there was almost never any overlap in either schizonts (Fig. 4Ai) which suggests that PfVps16Ks does not reside at the cis-Golgi. When looking at the trans-Golgi marker Rab6, some overlap could be seen with several PfVps16Ks foci whilst others had none (Fig. 4Aii). We next looked at the microneme markers AMA1 and EBA175 which are known to reside in different populations of these organelles(43-45). As seen with ERD2, minimal overlap was observed between the AMA1 and PfVps16Ks foci (Fig. 4Ai vs 4Aiii). However, a number of foci were very close/partially overlapping between EBA175 and PfVps16Ks though others were not colocalizing at all (Fig. 4Aiv). We finally looked at RAP1, a marker of the rhoptry organelle and saw that several foci were in very close juxtaposition whilst others seem to strongly overlap (Fig. 4Av). Quantification of the level of colocalization (Fig. 4B) revealed that PfVps16Ks overlapped more with Rab6 and RAP1 than ERD2 (Pearson’s correlation coefficient with Rab6: 0.44**±**0.02, RAP1: 0.44**±**0.04; Erd2: 0.27**±**0.03). AMA1 and EBA175 had coefficients in between with 0.35**±**0.04 and 0.38**±**0.06, respectively. The fact that not all foci share the same level of overlap suggests that PfVps16Ks is not a resident protein of either the trans-Golgi or the rhoptries but that it might perhaps shuttle between these organelles and/or others as seen with several proteins of the intracellular trafficking machinery such as PfSortilin(10, 11), PfVps45(46) and PfRbsn5L(47). The *T. gondii* components of the Vps-C complex TgVps11 and TgVps18 are critical for the generation of rhoptries and subsets of micronemes(26) and though this was not directly demonstrated for TgVps16, it is likely to be the case since it is part of the core complex(23). The partial colocalization of PfVps16Ks with the rhoptries and EBA175-containing micronemes suggest that the PfVps-C complex could potentially also be involved in their biogenesis, like its *T. gondii* counterpart.

**Fig. 4:**
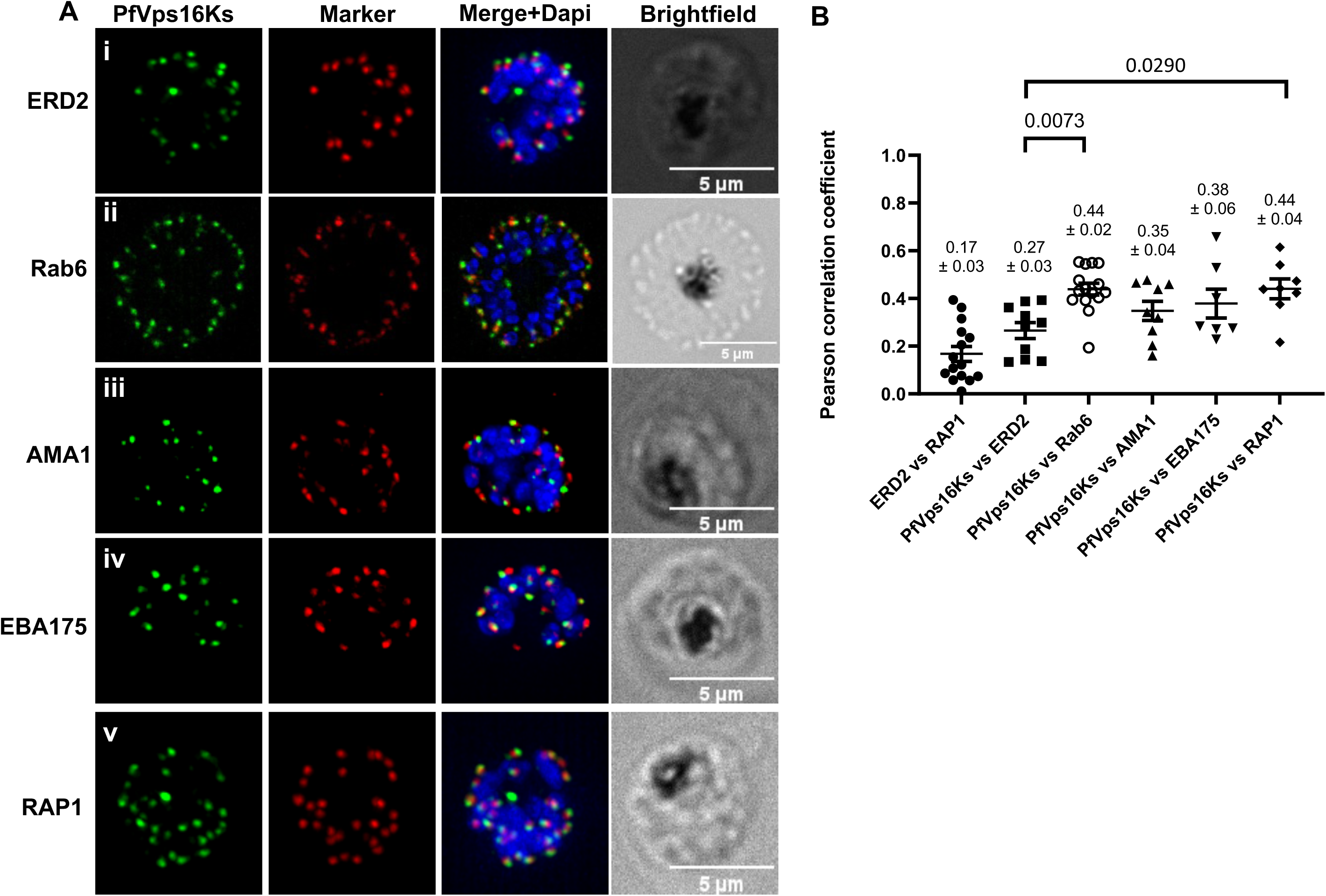
Colocalization analysis of PfVps16Ks. IFA on schizont stage parasites to determine the overlap between PfVps16Ks and: (Ai) the cis-Golgi marker ERD2; (Aii) the trans-Golgi marker Rab6; the micronemes markers (Aiii) AMA1 and (Aiv) EBA175; (Av) the rhoptry marker RAP1. (B) Pearson’s correlation analysis demonstrates that PfVps16 overlaps significantly more with Rab6 and RAP1 than with ERD2. ERD2 vs RAP1, n=15, used as a negative control. PfVps16Ks vs ERD2, n=10; PfVps16Ks vs Rab6, n=15; PfVps16Ks vs AMA1, n=9; PfVps16Ks vs EBA175, n=7; PfVps16Ks vs RAP1, n=8. Scale bar represents 5μm. Blue: DAPI stained nucleus. Values represent the mean ± standard error. *P-*values were calculated using one-way ANOVA followed by a Tukey’s multiple comparison test. Only the statistically significant p-values are shown.

### Attempts to mislocalize PfVps16Ks to the nucleus by knock sideways

To delve into the role of PfVps16 in the asexual blood stages, we performed Ks, a strategy that allows the conditional removal of a protein of interest from its site of action(48). To do so, we transfected the PfVps16Ks parasite line with a plasmid expressing a nuclear mislocalizer consisting of a triple nuclear localization signal fused to a double FKBP12-rapamycin binding domain and the mCherry fluorescent protein (3xNLS-2xFRB-cherry)(37). We first tested the ability of the mislocalizer to translocate PfVps16Ks to the nucleus when rapamycin was added to the medium (Rapa), thus removing it from its normal site of action. In the absence of Rapa, the mislocalizer overlapped with the DAPI-stained nucleus whilst PfVps16Ks showed its normal punctate pattern (Fig. 5Ai). When adding Rapa at the ring stage and letting the parasites mature to trophozoites and schizonts, GFP remained as foci that did not overlap with the DAPI (Fig. 5Aii and iii). Instead, some of the mCherry signal was now colocalizing with some of the GFP foci (Fig. 5Aii and iii, white arrows). This suggests that not only is PfVps16Ks not translocated into the nucleus but that some of the mislocalizer is actually delocalized from the nucleus by PfVps16Ks. This reverse mislocalization has previously been observed(49). We quantified this by calculating Pearson’s correlation coefficients. As seen in figures 4Bi and ii, there is no statistically significant difference in the overlap between DAPI and GFP or between GFP and mCherry with or without Rapa. However, the addition of Rapa leads to a decrease in the colocalization levels between DAPI and mCherry (Fig. 5Biii). These results might be potentially explained by a strong association of PfVps16Ks with the other members of the Vps-C complex preventing its extraction and subsequent translocation to the nucleus and resulting in some of the mislocalizer associating with the Vps-C complex. Perhaps the sandwich version of the Ks system using 2xFKBP both at the N- and C-terminus of GFP (2xFKBP-GFP-2xFKBP) would have been more efficient for nuclear mislocalization(37). Unfortunately, this lack of mislocalization prevented us from further investigating the role of PfVps16 in the biology of *P. falciparum*. Alternatives such as the DiCre-based(50) or TetR-Dozi-based(51) conditional expression systems are currently explored.

**Fig. 5:**
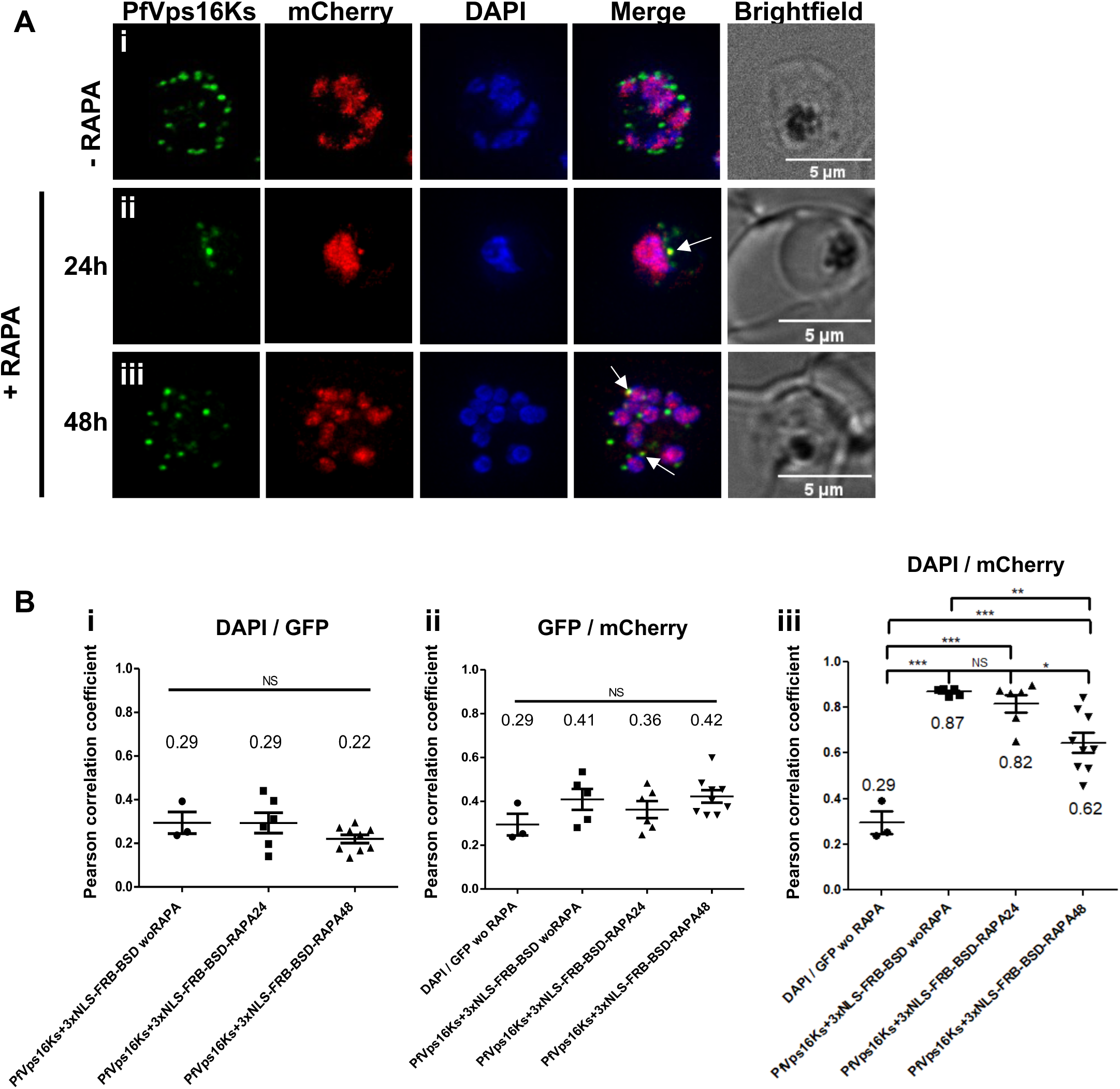
PfVps16Ks is not mislocalized to the nucleus upon addition of rapamycin. (A) Live microscopy showing that incubating the PfVps16Ks+mislocalizer parasite line with 250 nM rapamycin for 24h or 48h does not result in the delocalization of the green signal to the nucleus. Arrows show overlapping foci between PfVps16Ks and the mislocalizer potentially outside of the nucleus. Scale bar represents 5μm. Blue: DAPI stained nucleus. RAPA: rapamycin. (Bi) Pearson’s correlation analysis shows that the colocalization levels do not change between the DAPI and the PfVps16Ks (Bi) or the mCherry fused mislocalizer and PfVps16Ks (Bii). However, the colocalization between DAPI-stained nucleus and the mCherry-fused mislocalizer decreases upon the addition of rapamycin (Biii). Values represent the mean ± standard error. *P-*values were calculated using one-way ANOVA followed by a Tukey’s multiple comparison test. *: p<0.05; **: p<0.01. ***: p<0.001. NS=not significant. PfVps16Ks+3xNLS-FRB-BSD woRAPA, n=5. PfVps16Ks+3xNLS-FRB-BSD RAPA24, n=6. PfVps16Ks+3xNLS-FRB-BSD RAPA48, n=9.

In conclusion, our results suggests that PfVps16 and Vps33 have structural homology to their yeast and human counterparts and that their interaction are potentially conserved. Finally, partial colocalization of PfVps16 with markers of the trans-Golgi and the rhoptries suggests that the protein might cycle between the two organelles.

## MATERIALS AND METHODS

### Ethics statement

Study approved by the Canadian Blood Services (CBS) research ethics board, project number 2015.001 and by the CHU de Québec IRB, project number 2015–2230, B14-12-2230, SIRUL 104595. Written consent was obtained by the CBS for all study participants. Participants were informed about the study before providing consent. All experiments were performed in accordance with relevant guidelines and regulations.

#### Protein Structure Prediction

The protein sequence of PfVps16 (PF3D7_1239900) and PfVps33 (PF3D7_0935200) were retrieved from the *Plasmodium falciparum* genome database(58) (plasmodb.org). Their 3D structures were predicted using the AlphaFold server (https://alphafoldserver.com), which implements the AlphaFold 3 algorithm(27). All generated files are available in supplementary data.

#### Structural Alignment

The 3D structure of *Saccharomyces cerevisiae* Vps16 was isolated from the 3D structure of the Endosomal membrane tethering complex CORVET(24) (PDB: 8QX8). The 3D structure of human Vps33A was isolated from the Human Vps33A in complex with a fragment of human Vps16(33).To compare the predicted 3D structure of PfVps16 and PfVps33 with their homologous counterparts in yeast and humans respectively, we conducted a structural alignment analysis using US-align(59) (version 20240510) with default parameters (sequence independent structure alignment). TM-Score was used to quantify the degree of similarity between the two structures.

#### Figure Generation

All the figures of the 3D structures presented in this study were generated using the open-source software ChimeraX (version 1.7)(60).

### Parasite culture

*P. falciparum* NF54 asexual stage parasites were cultured under standard conditions in RPMI-HEPES medium at 4% hematocrit (human erythrocytes of O^+^ group) and 0.5% (w/v) Albumax^TM^ (Invitrogen) and kept at 37°C in a gas mixture of 5.0% oxygen, 5.0% carbon dioxide and 90% nitrogen(52).

#### Vector Construction and Transfection

To tag the endogenous PfVps16 with 2xFKBP-GFP, around 500 bp of the C-terminus of PfVps16 was amplified with primers 5’Not1-2571-PfVps16 (ATAgcggccgcGAATAGTCATATAAAATTTGTTCATACTTCC) and 3’AvrII-stopless-PfVps16 (ATAcctaggTCTTATGTTTGATATGGCATC) and cloned in frame with 2xFKBP-GFP in pSLI-2xFKBP-GFP-hDHFR digested NotI-AvrII(37). Parasites were transfected and integrants were selected as described previously with some modifications(37). Briefly, *P. falciparum* NF54 parasites were transfected with 100 µg of purified pSLI-PfVps16-2xFKBP-GFP plasmid. Positive selection for transfectants was achieved using 5nM WR99210 (WR). The drug resistant parasites were split into 3 separate wells with 2-4% parasitemia and went under another round of selection using 400µg/ml neomycin (NEO) to select for integrants. After parasite re-emergence (after around 10 days) WR was put back in the culture medium. Parasites were then cloned by limiting dilution resulting in the PfVps16Ks line. Genomic DNA was prepared from NEO and WR resistant parasites. Integration was monitored by PCR using the forward 5’upstream-2571-PfVps16-F (primer 1, GGGCAACATTCGCAAGC) and the reverse 3’90-GFP-R primer (primer 2) for 5’ integration and 5’pARL-F (primer 3) with the reverse 3’UTR-PfVps16-R primers (primer 4, GGTGAAAATAGAACTCGATGC) for the 3’ integration. Primer 1 was used with the reverse 3’-3’UTR-PfVps16-R primers (primer 4) to detect the WT version of the gene.

To generate the parasite line for the knock sideways, the PfVps16Ks line was transfected with 100 µg of purified p3xNLS-FRB-mCherry-BSD plasmid(37) and selected 2 µg/ ml blasticidin (Sigma-Aldrich) to obtain the PfVps16Ks+mislocalizer line.

### Western blotting

Saponin-extracted parasites from asynchronous PfVps16Ks line were harvested. Proteins were then separated on 10% (w/v) SDS-PAGE gel under reducing conditions and transferred to a PVDF membrane (Milipore). The membrane was blocked in 4% (w/v) milk in TBS-T. Antibodies used were mouse monoclonal anti-GFP 1:500 (ROCHE; IgG clone 7-1&13-1) and rabbit polyclonal anti-PfHSP70 1:40000 (StressMarq Bioscience Inc, SPC-186C)(53). Appropriate HRP-coupled secondary antibodies were used and immunoblots were developed using ECL (Biorad).

### Microscopy

Fluorescence microscopy acquisition was performed as previously described(41) using a GE Applied Precision Deltavision Elite microscope with a 100x 1.4NA objective and with a sCMOS camera and deconvolved with the SoftWorx software. For immunofluorescence assays, parasites were fixed with 4% paraformaldehyde-0.0075% glutaraldehyde (ProSciTech). After blocking in 3% bovine serum albumin (BSA fraction V, EMD), the slides were probed with combinations of antibodies: rabbit anti-ERD2 (MRA-72, 1:1000)(40); rabbit anti-AMA1, 1:1000(54) or mouse monoclonal anti-AMA1 (clone 1F9, 1:500)(55); rabbit anti-PfEBA175 (1:1000)(56), mouse anti-RAP1 (1:3000)(57). Primary antibodies were probed with Alexa Fluor 594 anti-rabbit IgG or anti-mouse IgG (Molecular Probes) and Alexa Fluor 488 anti-rabbit IgG or anti-mouse IgG (Cell Signaling). Slides were mounted with 4’,6-Diamidino-2-phenylindole dihydrochloride (DAPI, Invitrogen, 100ng/ul) in VectaShield (Vector Labs) or ProLong Gold anti-fade (Molecular Probes). Pearson’s correlation coefficient between Alexa488 and Alexa594 channels was calculated on deconvolved regions of interests of image stacks, including zero-zero pixels and without thresholding using the SoftWorx software (GE). Data were analyzed for statistical significance using one-way ANOVA followed by a Tukey multiple comparison test. Chromatic calibration of the microscope was performed prior to imaging experiments.

#### Knock sideways attempts

Tightly synchronous ring stage PfVps16Ks+mislocalizer parasites were seeded at 2% parasitemia and grown **+/-** 250 nM rapamycin. After 24h and 48h in culture, the cells were harvested, stained with 4’,6-Diamidino-2-phenylindole dihydrochloride (DAPI, Invitrogen, 100ng/ul) and imaged immediately.

#### Statistical analysis

Prism 7(GraphPad) was used for all statistical analyses. Depending on the assay, one-way ANOVA or two-tailed unpaired t-tests were performed. A *P* value of < 0.05 was considered statistically significant.

## Materials and Correspondence

Requests for materials should be addressed to Dave Richard.

## Acknowledgments

We would like to thank Tobias Spielman for the pSLI plasmids and Robin Anders, Michael Blackman and Alan Cowman for antibodies. We also thank Jacobus Pharmaceuticals for WR99210. The following reagents were obtained through MR4 as part of the BEI Resources, National Institute of Allergy and Infectious Diseases, National Institutes of Health, USA: Polyclonal Anti-Plasmodium falciparum PfERD2 (antiserum, Rabbit) and *Plasmodium falciparum*, Strain NF54 (Patient Line E), MRA-1000, contributed by Megan G. Dowler. We would also like to acknowledge the Canadian Blood Services for providing human erythrocytes. The authors declare no competing financial interests. This study was funded through a National Science and Engineering Council of Canada (RGPIN-2018-06281). DR is a Fonds de la Recherche du Québec-Santé Senior fellow.

## Author contributions

T.G. performed most of the experimental work, interpreted results and edited the manuscript. F.L. performed the structural analyses, interpreted results and edited the manuscript. A.S. performed the imaging and analysis of PfVps16Ks and Rab6, interpreted results and edited the manuscript. D.R. conceived the study, designed experiments, interpreted results and wrote the manuscript.

## Data availability

The data supporting the findings of this study are available within the paper and are also available from the corresponding author upon request.

## Figure titles

**Figure S1:**
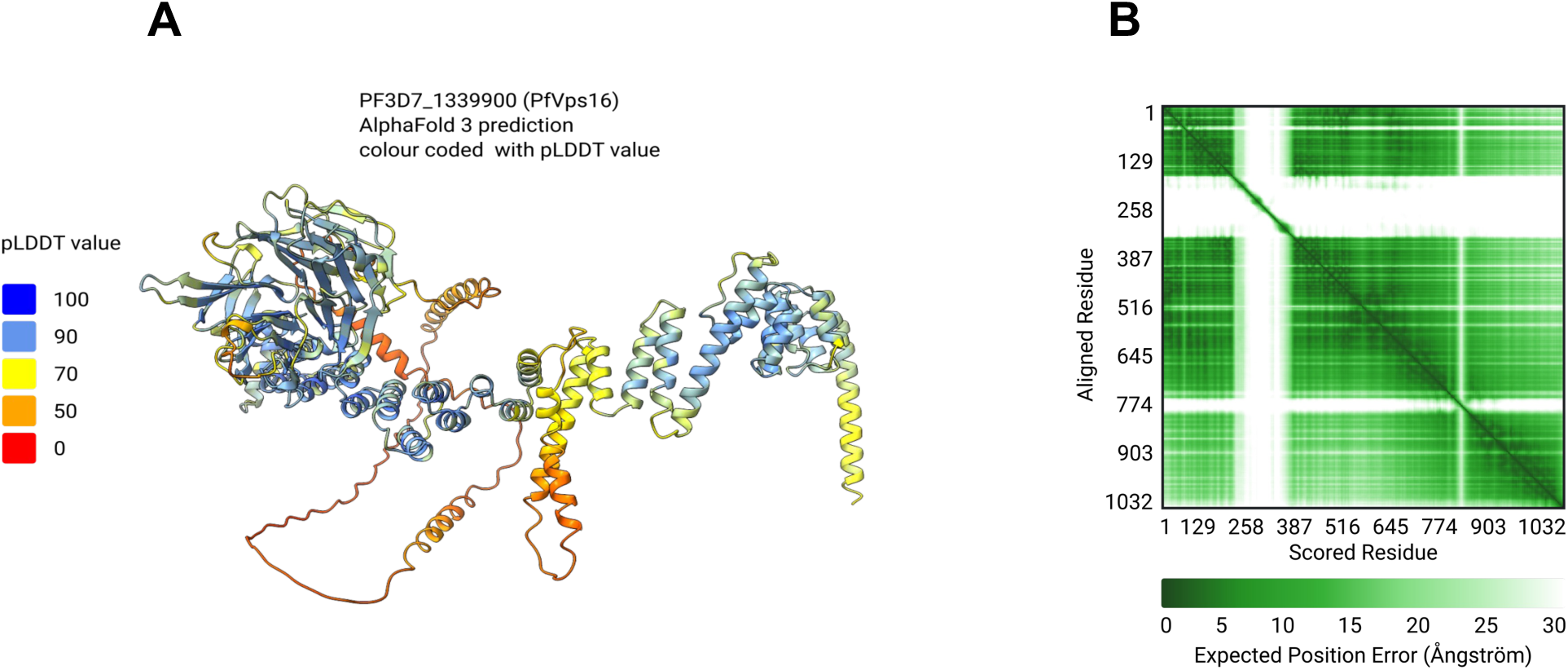
AlphaFold 3 structure prediction of PfVps16. (A) PfVps16 3D structure color-coded with predicted local-distance difference test (pLDDT) value. Predicted template modeling (pTM) score = 0.68. (B) Plot of the predicted aligned error (PAE).

**Figure S2:**
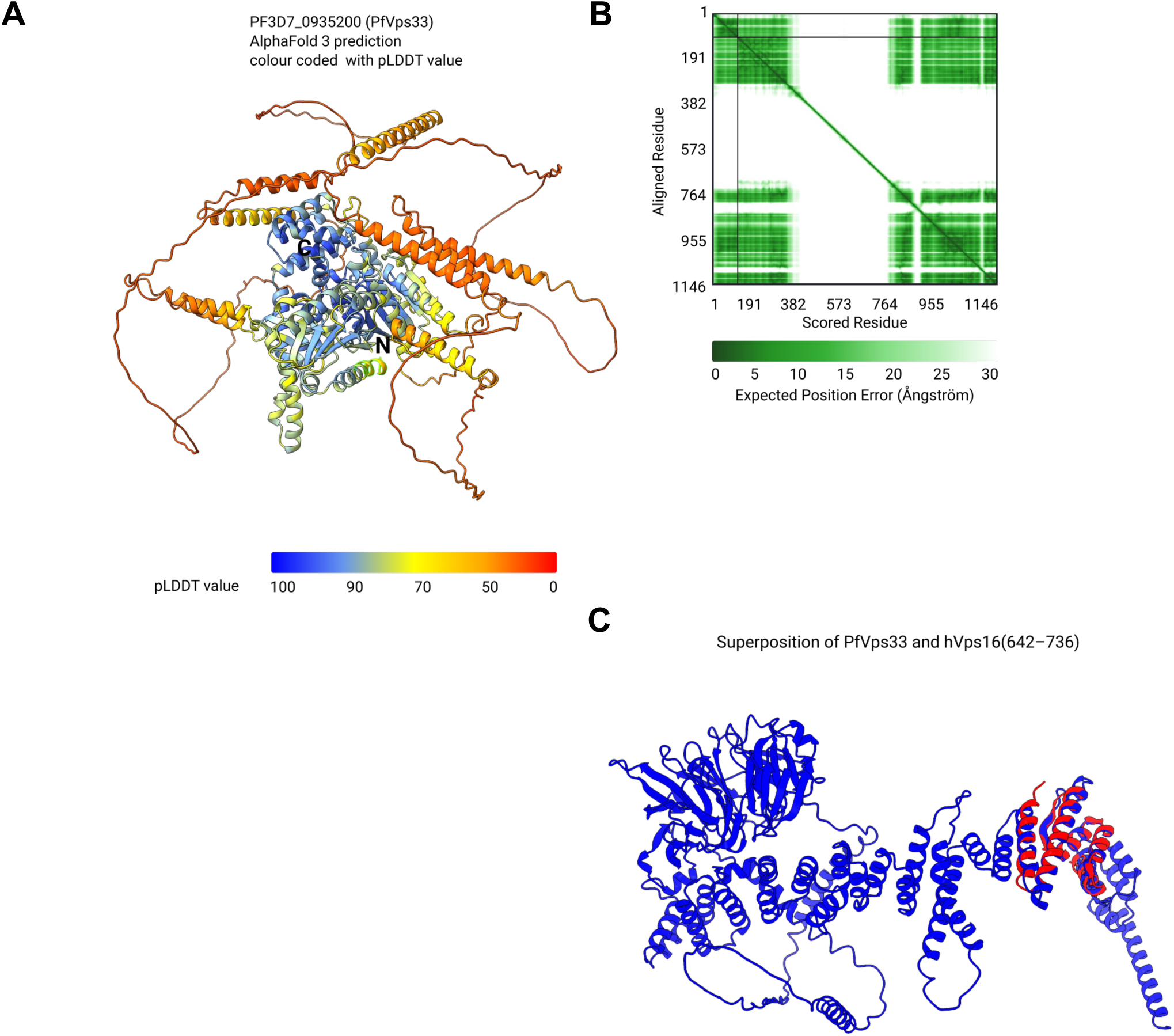
AlphaFold 3 structure prediction of PfVps33 and comparison with hVps33. (A) PfVps33 3D structure colour-coded with predicted local-distance difference test (pLDDT) value. Predicted template modeling (pTM) score = 0.58. (B) Plot of the predicted aligned error (PAE). (C) Superposition of PfVps33 in blue with hVps16 (residues 642-736) in red.

